# A complete time-calibrated multi-gene phylogeny of the European butterflies

**DOI:** 10.1101/844175

**Authors:** Martin Wiemers, Nicolas Chazot, Christopher W. Wheat, Oliver Schweiger, Niklas Wahlberg

## Abstract

With the aim of supporting ecological analyses in butterflies, the third most species-rich superfamily of Lepidoptera, this paper presents the first time-calibrated phylogeny of all 496 extant butterfly species in Europe, including 18 very localized endemics for which no public DNA sequences had been available previously. It is based on a concatenated alignment of the mitochondrial gene COI and up to 11 nuclear gene fragments, using Bayesian inference of phylogeny. To avoid analytical biases that could result from our region-focus sampling, our European tree was grafted upon a global genus-level backbone butterfly phylogeny for analyses. In addition to a consensus tree, we provide the posterior distribution of trees and the fully-concatenated alignment for future analyses.

## Background & Summary

The incorporation of phylogenetic information in ecological theory and research has led to significant advancements by facilitating the connection of large-scale and long-term macro-evolutionary processes with ecological processes in the analysis of species interactions with their abiotic and biotic environments^1,2^. Phylogenies are increasingly used across diverse areas of macroecological research^3^, such as studies on large-scale diversity patterns^4^, disentangling historical and contemporary processes^5^, latitudinal diversity gradients^6^ or improving species area relationships^7^. Phylogenetic information has also improved studies on assembly rules of local communities^8–10^, including spatiotemporal community dynamics^11^ and multi-spatial and -temporal context-dependencies^12^. Additionally, phylogenetic information has provided insights into the mechanisms and consequences of biological invasions^13–16^. They also contribute to assessments of ecosystem functioning and service provisioning^17,18^, though phylogenetic relationships cannot simply be taken as a one-to-one proxy for ecosystem functioning^19,20^. However, they are of great value for studies of species traits and niche characteristics by quantifying the amount of phylogenetic conservatism^21^ and ensuring statistical independence^22^ in multi-species studies. Using an ever increasing toolkit of phylogenetic metrics^23,24^, and a growing body of phylogenetic insights, the afore mentioned advances across diverse research fields document how integrating evolutionary and ecological information can enhance assessments of future impacts of global change on biodiversity^25–27^ and consequently inform conservation efforts (but see also ^20,28^).

Although the amount of molecular data has increased exponentially during the last decades, most available phylogenetic studies are either restricted to a selected subset of species, higher taxa, or to small geographic areas. Complete and dated species-level phylogenetic hypotheses for species-rich taxa of larger regions are usually restricted to vascular plants^29^ or vertebrates, such as global birds^30^ or European tetrapods^31^, or the analyses are based on molecular data from a small subset of species (e.g. 5% in ants^6^). Surprisingly, comparable phylogenetic hypotheses are rare for insects, which comprise the majority of multicellular life on earth^32^, have enormous impacts on ecosystem functioning, provide a multitude of ecosystem services^33^, and have long been used as biodiversity indicators^34^.

Here, we present the first comprehensive time-calibrated molecular phylogeny of all 496 extant European butterfly species (Lepidoptera: Papilionoidea), based on one mitochondrial and up to eleven nuclear genes, and the most recent systematic list of European butterflies^35^. European butterflies are well-studied, ranging from population level analyses^36^ to large-scale impacts of global change^37^, with good knowledge on species traits and environmental niche characteristics^38,39^, population trends^40,41^ and large-scale distributions^42,43^ and are thus well placed for studies in the emerging field of ecophylogenetics^1^.

Compared to other groups of insects, the phylogenetic relationships of butterflies are reasonably well-known, with robust backbone molecular phylogenies at the subfamily^44–46^ and genus-level^47^. In addition, molecular phylogenies also exist for most butterfly families^48–58^ as well as major subgroups^59–65^ and comprehensive COI data on species level are available from DNA barcoding studies^66–71^. Some ecological studies on butterflies have already incorporated phylogenetic information, e.g. on the impact of climate change on abundance trends^72,73^, the sensitivity of butterflies to invasive species^13,74^ or the ecological determinants of butterfly vulnerability^75^. However, the phylogenetic hypotheses used in these studies had incomplete taxon coverage (but see ^76^) and were not made available for reuse by other researchers. To fill these gaps in the literature, and to facilitate the growing field of ecophylogenetics, here we present the first complete and time-calibrated species-level phylogeny of a speciose higher invertebrate taxon above the family level for an entire continent. Importantly, we provide this continent-wide fully resolved phylogeny in standard analysis formats for further advancements in theoretical and applied ecology.

## Methods

### Taxonomic, spatial and temporal coverage

We analyse a dataset comprising all extant European species of butterflies (Papilionoidea), including the families Papilionidae, Hesperiidae, Pieridae, Lycaenidae, Riodinidae and Nymphalidae. We base our species concepts, as well as the area defined as Europe, on the latest checklist of European butterflies^35^.

### Acquisition of sequence data

The data were mainly collated from published sources and downloaded from NCBI GenBank (Table S1). One mitochondrial gene, cytochrome c oxidase subunit I (COI, 1464 bp), was available for all species in the data matrix, in particular the 5’ half of the gene (658 bp, also known as the DNA barcode). Eleven nuclear genes were included when available: elongation factor-1α (EF-1α, 1240 bp), ribosomal protein S5 (RpS5, 617 bp), ribosomal protein S2 (RpS2, 411 bp), carbamoylphosphate synthase domain protein (CAD, 850 bp), cytosolic malate dehydrogenase (MDH, 733 bp), glyceraldehyde-3-phosphate dehydrogenase (GAPDH, 691 bp), isocitrate dehydrogenase (IDH, 711 bp), wingless (412 bp), Arginine kinase (ArgK, 596 bp) and Dopa Decarboxylase (DDC, 373 bp) and histone 3 (H3, 329 bp). H3 has been sequenced almost exclusively from the family Lycaenidae, while the other gene regions have been sampled widely also in the other butterfly families. For each gene, the longest available sequence was used. However, in the case of several available sequences of similar length, those of European origin were preferentially used. Sequences were aligned manually to maintain protein reading frame, and were curated and managed using VoSeq^77^.

In several cases, new sequences were generated for this study. For these specimens, protocols followed Wahlberg and Wheat ^78^ or Wiemers and Fiedler ^66^. These include several species that did not have any available published sequences, many of which are island endemics (Table 1). The new sequences have been submitted to GenBank (accessions xxxx-xxxx).

**Table 1.** Newly sequenced species for which no published sequences had previously been available

| **Taxon** | **Origin** |
| --- | --- |
| *Coenonympha orientalis* | Greece |
| *Glaucopsyche paphos* | Cyprus |
| *Gonepteryx maderensis* | Portugal: Madeira |
| *Hipparchia azorina* | Portugal: Azores |
| *Hipparchia bacchus* | Spain: Canary Islands |
| *Hipparchia cretica* | Greece: Crete |
| *Hipparchia gomera* | Spain: Canary Islands |
| *Hipparchia maderensis* | Portugal: Madeira |
| *Hipparchia mersina* | Greece: Samos |
| *Hipparchia miguelensis* | Portugal: Madeira |
| *Hipparchia sbordonii* | Italy: Pontine Islands |
| *Hipparchia tamadabae* | Spain: Canary Islands |
| *Hipparchia tilosi* | Spain: Canary Islands |
| *Hipparchia wyssii* | Spain: Canary Islands |
| *Lycaena bleusei* | Spain |
| *Pieris balcana* | North Macedonia |
| *Pieris wollastoni* | Portugal: Madeira |
| *Thymelicus christi* | Spain: Canary Islands |

Almost all genera are represented by multiple genes, except *Borbo, Gegenes, Laeosopis, Callophrys* and *Cyclyrius* (the latter recently synonymized with *Leptotes*^79^) which are represented only by the COI gene. Species represented by only the DNA barcode tend to be closely related to species with more genes sequenced (Table S1), minimizing the potential bias these samples could have in our analyses.

### Phylogenetic tree reconstruction

A biogeographically restricted tree of a given taxon is inherently very asymmetrically sampled. To avoid potentially strong biases when estimating topology and divergence times we chose to build upon the recent genus-level tree of butterflies^47^, which provides a well-supported time-calibrated backbone and corresponds well with a recent phylogenomic analysis of Lepidoptera^80^. This backbone tree contains 994 taxa, each taxon representing a genus across all Papilionoidea. The tree was time-calibrated using a set of 14 fossil calibration points, which provided minimum ages and 10 calibration points based on ages of host plant clades taken from the literature, which provided maximum ages. Importantly, Chazot, et al. ^47^ tested the robustness of their results to a wide range of alternative assumptions made in the time-calibration analysis, and showed that the estimated times of divergences were robust.

### Analysis overview

In order to produce our time-calibrated tree of European butterflies, we identified the position of the European lineages and designed a grafting procedure accordingly. We split the European butterflies that needed to be added to the tree into 12 subclades. For each of these subclades we combined the DNA sequences of the taxa already included in the backbone to the DNA sequences of the European taxa to assemble an aligned molecular matrix. After identifying the best partitioning scheme, we performed a tree reconstruction without time-calibration (only estimating relative branch lengths). The subclade trees were then rescaled using the ages estimated in the backbone and were subsequently grafted. This procedure was repeated using 1000 trees from BEAST posterior distributions of the backbone and subclade trees in order to obtain a posterior distribution of grafted trees. The details of these procedures are described below.

### Backbone and subclades

The time-calibrated backbone tree provided by Chazot, et al. ^47^ contained about 55% of all butterfly genera, including most genera occurring in Europe. A fixed topology was obtained using RAxML^81^ and node ages where estimated with BEAST v.1.8.3.^82^. We used this fixed topology from Chazot, et al. ^47^ to identify at which nodes European clades should be grafted. We partitioned the analysis into 12 subclades. For each subclade, the DNA sequences of all taxa already included in the global backbone (including also non-European taxa) were combined with the DNA sequences of all the new European taxa that were added. In addition to the focal taxa, we added between two and four outgroups.

The subclades, sorted by families, were defined as follows:

Papilionidae – All Papilionidae were placed into one subclade.

Hesperiidae – We identified two main clades to graft within the Hesperiidae: Hesperiinae and Pyrginae. The Hesperiinae subclade was extended to also encompass the subfamilies Heteropterinae and Trapezitinae. The genus *Muschampia*, not available in the backbone, was included in the Pyrginae subclade.

Pieridae – All Pieridae were considered as a single clade.

Lycaenidae – All Lycaenidae were considered as a single clade.

Riodinidae – The only European Riodinidae species, *Hamearis lucina*, was already available in the backbone tree.

Nymphalidae – European Nymphalidae were divided into seven subclades. (i) A subclade for the Apaturinae. (ii) In order to add *Danaus chrysippus* we generated a tree of Danainae. (iii) We combined the sister clades Heliconiinae and Limenitidinae into a single subclade. (iv) Nymphalinae was treated as a single subclade. (v) A first clade of Satyrinae contained the genera *Kirinia*, *Pararge*, *Lasiommata*, *Tatinga*, *Chonala* and *Lopinga*. (vi) A second Satyrinae clade contained the genera *Calisto, Euptychia, Callerebia, Proterebia, Gyrocheilus, Strabena, Ypthima, Ypthimomorpha*, *Stygionympha*, *Cassionympha*, *Neocoenyra*, *Pseudonympha*, *Erebia*, *Boerebia*, *Hyponephele*, *Cercyonis*, *Maniola*, *Aphantopus*, *Pyronia*, *Faunula*, *Grumia*, *Paralasa*, *Melanargia*, *Hipparchia*, *Berberia*, *Oeneis*, *Neominois*, *Karanasa*, *Brintesia*, *Arethusana*, *Satyrus*, *Pseudochazara* and *Chazara*. (vii) A third Satyrinae clade was created for the genus *Coenonympha*. Charaxinae were not treated separately from the backbone. *Charaxes jasius* is the only Charaxinae occuring in Europe and *Charaxes castor* (which is very closely related to *C. jasius*^83^) was already included in the backbone tree from Chazot et al. ^47^. Hence, we used the position of *Charaxes castor* for *Charaxes jasius*.

### Partitioning the dataset

For each subclade we ran Partition Finder 2 ^84^ in order to partition the data and choose substitution models. The dataset was initially partitioned into genes and codon positions. Branch lengths were set to *linked* and the comparison between partitioning strategies was made using the greedy algorithm and BIC score^85^.

### Phylogenetic reconstruction

For each subclade, the dataset was imported in BEAUTi v.1.8.3^86^ and partitioned according to the partitioning strategy identified by Partition Finder 2. We enforced the monophyly of the clade to be grafted (i.e., excluding the outgroups). All other relationships were estimated by BEAST v.1.8.3.^82^. We used an uncorrelated relaxed clock with lognormal distribution. By default, we started by setting one molecular clock per partition. If convergence or good mixing could not be obtained after running BEAST we reduced the number of molecular clocks (see details for each dataset further below). We did not add any time-calibration and therefore only estimated the relative timing of divergence. We performed at least two independent runs with BEAST for each subclade. We checked for convergence and mixing of the MCMC using Tracer v.1.6.0^87^ and in the case of full convergence of the runs, the posterior distribution of trees from different runs were combined after removing the burn-in fraction.

### Grafting procedure

Subclades were grafted on the backbone as follows. One backbone was sampled from the posterior distribution of time-calibrated trees from Chazot, et al.^47^. For each subclade, one subclade tree was sampled from the posterior distribution of trees, the outgroups removed and the tree was rescaled based on the crown age of the subclade extracted from the backbone tree. Finally, the rescaled subclade tree was grafted on the backbone after removing all lineages belonging to this subclade in the backbone (i.e. only keeping the stem branch). We repeated this procedure for 1000 backbone trees and 1000 subclade trees and thus we obtained a posterior distribution of 1000 grafted trees. The topology of the backbone was fixed (see ^47^) but the topologies of the subclades were free. Hence the posterior distribution of grafted trees includes a posterior distribution of topologies and node ages.

We describe below the details of the phylogenetic tree reconstruction for each subclade.

#### 1- Papilionidae

*Dataset* – The dataset for the Papilionidae consisted of 36 taxa to which three outgroups were added: *Macrosoma tipulata* (Hedylidae), *Achlyodes busiris* (Hesperiidae), *Pieris rapae* (Pieridae). We concatenated 11 gene fragments (COI, CAD, EF-1α, GAPDH, ArgK, IDH, MDH, RpS2, RpS5, DDC, wingless).

*Partition Finder* – Partition Finder identified 12 subsets.

| Subset # | Substitution model | Gene fragments and codon positions |
| --- | --- | --- |
| Subset 1 | TRN+I+G | ArgK_pos1, RpS2_pos1, EF-1α_pos1, RpS5_pos1 |
| Subset 2 | TRN+I+G | COI-begin_pos2, RpS5_pos2, ArgK_pos2, GAPDH_pos2, CAD_pos2, MDH_pos2, IDH_pos2, COI-end_pos2 |
| Subset 3 | HKY | ArgK_pos3 |
| Subset 4 | HKY+I+G | CAD_pos3 |
| Subset 5 | GTR+G | wingless_pos1, IDH_pos1, GAPDH_pos1, MDH_pos1, CAD_pos1 |
| Subset 6 | TRN+G | COI-end_pos3, COI-begin_pos3 |
| Subset 7 | GTR+G | DDC_pos2, COI-end_pos1, COI-begin_pos1 |
| Subset 8 | GTR+I+G | DDC_pos3, wingless_pos3, RpS2_pos3 |
| Subset 9 | K80+G | DDC_pos1, wingless_pos2 |
| Subset 10 | GTR+G | EF-1α_pos3 |
| Subset 11 | TRNEF+I | RpS2_pos2, EF-1α_pos2 |
| Subset 12 | GTR+G | RpS5_pos3, GAPDH_pos3, IDH_pos3, MDH_pos3 |

*BEAST analysis* – In order to improve the quality of our runs we replaced the default priors for rates of substitutions by uniform prior ranging between 0 and 10 for the following cases: subset5.at, subset5.cg, subset7.cg, subset7.gt, subset12.cg, subset12.gt. We used one molecular clock per subset identified by Partition Finder and obtained good mixing and convergence. We used a Birth-Death tree prior. We performed three runs of 40 million generations, sampling trees and parameters every 4000 generations.

*Grafting* – For grafting, the outgroups were removed, as well as *Baronia brevicornis*, the first Papilionidae to diverge and endemic to Mexico, i.e. we grafted at the most recent common ancestor (mrca) of all Papilionidae but *Baronia brevicornis*.

#### 2- Hesperiidae: Hesperiinae

*Dataset* – The dataset for the Hesperiinae consisted of 169 taxa to which two outgroups were added: *Typhedanus ampyx* (Hesperiidae), *Mylon pelopidas (*Hesperiidae). We concatenated 10 gene fragments (COI, CAD, EF-1α, GAPDH, ArgK, IDH, MDH, RpS2, RpS5, wingless).

*Partition Finder* – Partition Finder identified 17 subsets.

| Subset # | Substitution model | Gene fragments and codon positions |
| --- | --- | --- |
| Subset 1 | GTR+I+G | COI-end_pos1, COI-begin_pos1, ArgK_pos1 |
| Subset 2 | GTR+I+G | ArgK_pos2, IDH_pos2, EF-1α_pos2 |
| Subset 3 | GTR+G | ArgK_pos3 |
| Subset 4 | TRN+I+G | CAD_pos3, IDH_pos3 |
| Subset 5 | GTR+I+G | IDH_pos1, CAD_pos1 |
| Subset 6 | GTR+I+G | CAD_pos2, MDH_pos2 |
| Subset 7 | TRN+I+G | COI-begin_pos3, COI-end_pos3 |
| Subset 8 | HKY+I+G | RpS5_pos2, COI-begin_pos2, GAPDH_pos2, COI-end_pos2 |
| Subset 9 | GTR+I+G | EF-1α_pos3 |
| Subset 10 | GTR+I+G | EF-1α_pos1, MDH_pos1 |
| Subset 11 | GTR+I+G | RpS2_pos3, GAPDH_pos3, RpS5_pos3 |
| Subset 12 | GTR+I+G | GAPDH_pos1, wingless_pos2 |
| Subset 13 | TRN+I+G | MDH_pos3 |
| Subset 14 | JC+I | RpS2_pos2 |
| Subset 15 | GTR+I+G | RpS2_pos1, RpS5_pos1 |
| Subset 16 | GTR+I+G | wingless_pos3 |
| Subset 17 | SYM+I+G | wingless_pos1 |

*BEAST analysis* – Preliminary analyses showed problems with the subset 3 (ArgKin_pos3) and was therefore removed from the analyses. In order to improve the quality of our runs we replaced the default priors for rates of substitutions by uniform priors ranging between 0 and 10 for the following cases: subset17.cg. The substitution model for the subset 14 was also changed into HKY+I after preliminary analyses. We used one molecular clock per subset identified by Partition Finder and obtained good mixing and convergence. We used a Birth-Death tree prior. We performed two runs of 150 million generations, sampling trees and parameters every 15000 generations.

*Grafting* – For grafting, the outgroups were removed and the subclade grafted at the mrca of Hesperiinae.

#### 3- Hesperiidae: Pyrginae

*Dataset* – The dataset for the Pyginae consisted of 77 taxa to which three outgroups were added: *Typhedanus ampyx* (Hesperiidae), *Pyrrhopyge zenodorus (*Hesperiidae) and *Hasora khoda* (Hesperiidae). We concatenated 10 gene fragments (COI, CAD, EF-1α, GAPDH, ArgK, IDH, MDH, RpS2, RpS5, wingless).

*Partition Finder* – Partition Finder identified 14 subsets.

| Subset # | Substitution model | Gene fragments and codon positions |
| --- | --- | --- |
| Subset 1 | TRN+I+G | EF-1α_pos1, IDH_pos1, RpS2_pos1, RpS5_pos1, ArgK_pos1 |
| Subset 2 | TRN+I+G | EF-1α_pos2, GAPDH_pos2, MDH_pos2, COI-begin_pos2, COI-end_pos2, IDH_pos2, CAD_pos2, RpS5_pos2, ArgK_pos2 |
| Subset 3 | GTR+G | ArgK_pos3 |
| Subset 4 | HKY+I+G | CAD_pos3 |
| Subset 5 | GTR+I+G | CAD_pos1, MDH_pos1, GAPDH_pos1 |
| Subset 6 | HKY+G | COI-begin_pos3 |
| Subset 7 | GTR+I+G | COI-end_pos1, COI-begin_pos1 |
| Subset 8 | TRN+I+G | COI-end_pos3 |
| Subset 9 | GTR+G | EF-1α_pos3 |
| Subset 10 | GTR+I+G | RpS2_pos3, GAPDH_pos3, RpS5_pos3 |
| Subset 11 | GTR+I+G | MDH_pos3, IDH_pos3 |
| Subset 12 | JC+G | wingless_pos2, RpS2_pos2 |
| Subset 13 | HKY+G | wingless_pos3 |
| Subset 14 | SYM+G | wingless_pos1 |

*BEAST analysis* – In order to improve the quality of our runs we replaced the default priors for rates of substitutions by uniform prior ranging between 0 and 10 for the following cases: subset7.ac, subset7.gt, subset14.cg, subset3.cg. Preliminary analyses showed problems when using a separate molecular clock for each subset identified by Partition Finder. We restricted the analysis to one molecular clock. We used a Birth-Death tree prior. We performed two runs of 100 million generations, sampling trees and parameters every 10000 generations.

*Grafting* – For grafting, the outgroups were removed and the subclade grafted at the mrca of Pyrginae.

#### 4- Pieridae

*Dataset* – The dataset for the Pieridae consisted of 126 taxa to which three outgroups were added: *Bicyclus anynana* (Nymphalidae), *Achylodes busiris (*Hesperiidae) and *Papilio glaucus* (Papilionidae). We concatenated 11 gene fragments (COI, CAD, EF-1α, GAPDH, ArgK, IDH, MDH, RpS2, RpS5, DDC, wingless).

*Partition Finder* – Partition Finder identified 17 subsets.

| Subset # | Substitution model | Gene fragments and codon positions |
| --- | --- | --- |
| Subset 1 | GTR+I+G | ArgK_pos1, EF-1α_pos1, GAPDH_pos1 |
| Subset 2 | TRN+I+G | ArgK_pos2, DDC_pos2, CAD_pos2, IDH_pos2 |
| Subset 3 | KHY | ArgK_pos3 |
| Subset 4 | HKY+I+G | CAD_pos3 |
| Subset 5 | GTR+I+G | CAD_pos1, RpS2_pos1, IDH_pos1, RpS5_pos1, MDH_pos1 |
| Subset 6 | HKY+G | COI-begin_pos3 |
| Subset 7 | GTR+I+G | COI-begin_pos1, COI-end_pos1 |
| Subset 8 | GTR+I+G | MDH_pos2, COI-begin_pos2, RpS5_pos2, COI-end_pos2, EF-1α_pos2, GAPDH_pos2 |
| Subset 9 | TRN+I+G | COI-end_pos3 |
| Subset 10 | K80+G | DDC_pos3, RpS2_pos3 |
| Subset 11 | SYM+G | DDC_pos1, wingless_pos1 |
| Subset 12 | GTR+I+G | EF-1α_pos3 |
| Subset 13 | GTR+I+G | RpS5_pos3, GAPDH_pos3 |
| Subset 14 | GTR+I+G | IDH_pos3, MDH_pos3 |
| Subset 15 | JC+I | RpS2_pos2 |
| Subset 16 | GTR+I+G | wingless_pos3 |
| Subset 17 | K80+G | wingless_pos2 |

*BEAST analysis* – In order to improve the quality of our runs we replaced the default priors for rates of substitutions by uniform prior ranging between 0 and 10 for the following case: subset7.cg. The substitution model for the subset 7 was also changed into GTR+G after preliminary analyses. We used one molecular clock per subset identified by Partition Finder and obtained good mixing and convergence. We used a Birth-Death tree prior. We performed two runs of 100 million generations, sampling trees and parameters every 10000 generations.

*Grafting* – For grafting, the outgroups were removed and the subclade grafted at the mrca of Pieridae.

#### 5- Lycaenidae

*Dataset* – The dataset for the Lycaenidae consisted of 187 taxa to which three outgroups were added: *Bicyclus anynana* (Nymphalidae), *Pieris rapae (*Pieridae) and *Hamearis lucina* (Riodinidae). We concatenated 12 gene fragments (COI, CAD, EF-1α, GAPDH, ArgK, IDH, MDH, RpS2, RpS5, DDC, wingless and H3).

*Partition Finder* – Partition Finder identified 12 subsets.

| Subset # | Substitution model | Gene fragments and codon positions |
| --- | --- | --- |
| Subset 1 | TRN+I+G | ArgK_pos1, H3_pos1, EF-1α_pos1, RpS5_pos1, GAPDH_pos1 |
| Subset 2 | GTR+I+G | RpS5_pos2, GAPDH_pos2, ArgK_pos2, DDC_pos2, MDH_pos2, COI-begin_pos2, COI-end_pos2, EF-1α_pos2, IDH_pos2, CAD_pos2 |
| Subset 3 | GTR+G | H3_pos3, ArgK_pos3, wingless_pos3 |
| Subset 4 | HKY+I+G | CAD_pos3 |
| Subset 5 | GTR+G | wingless_pos2, RpS2_pos1, MDH_pos1, IDH_pos1, CAD_pos1 |
| Subset 6 | GTR+G | COI-begin_pos3, COI-end_pos3 |
| Subset 7 | GTR+I+G | COI-begin_pos1, COI-end_pos1 |
| Subset 8 | TRNEF | DDC_pos3, RpS2_pos3 |
| Subset 9 | SYM+G | wingless_pos1, DDC_pos1 |
| Subset 10 | GTR+I+G | EF-1α_pos3 |
| Subset 11 | GTR+G | MDH_pos3, RpS5_pos3, GAPDH_pos3, IDH_pos3 |
| Subset 12 | JC+I | RpS2_pos2, H3_pos2 |

*BEAST analysis* – In order to improve the quality of our runs we replaced the default priors for rates of substitutions by uniform prior ranging between 0 and 10 for the following cases: subset3.cg, subset6.ag, subset6.at, subset11.gt_subst7.cg. We used one molecular clock per subset identified by Partition Finder and obtained good mixing and convergence. We used a Birth-Death tree prior. We performed two runs of 150 million generations, sampling trees and parameters every 15000 generations.

*Grafting* – For grafting, the outgroups were removed and the subclade grafted at the mrca of Lycaenidae.

#### 6- Nymphalidae: Danainae

*Dataset* – The dataset for the Danainae consisted of 7 taxa to which two outgroups were added: *Euploea camaralzeman* (Nymphalidae) and *Lycorea halia (*Nymphalidae). We concatenated 9 gene fragments (COI, CAD, EF-1α, GAPDH, IDH, MDH, RpS2, RpS5, wingless).

*Partition Finder* – Partition Finder identified 8 subsets.

| Subset # | Substitution model | Gene fragments and codon positions |
| --- | --- | --- |
| Subset 1 | HKY+I | MDH_pos3, CAD_pos3 |
| Subset 2 | GTR+G | RpS5_pos1, RpS2_pos1, GAPDH_pos1, EF-1α_pos1, wingless_pos1, MDH_pos1, CAD_pos1, IDH_pos1 |
| Subset 3 | HKY+I | COI-end_pos2, COI-begin_pos2, RpS5_pos2, GAPDH_pos2, MDH_pos2, EF-1α_pos2, CAD_pos2, IDH_pos2 |
| Subset 4 | HKY+G | COI-end_pos3, COI-begin_pos3 |
| Subset 5 | TRN+G | COI-end_pos1, COI-begin_pos1 |
| Subset 6 | HKY+G | EF-1α_pos3, RpS5_pos3 |
| Subset 7 | GTR+G | GAPDH_pos3, RpS2_pos3, IDH_pos3, wingless_pos3 |
| Subset 8 | JC | RpS2_pos2, wingless_pos2 |

*BEAST analysis* – We used one molecular clock per subset identified by Partition Finder and obtained good mixing and convergence. We used a Birth-Death tree prior. We performed two runs of 20 million generations, sampling trees and parameters every 2000 generations.

*Grafting* – For grafting, the outgroups were removed and the subclade grafted at the mrca of Danainae.

#### 7- Nymphalidae: Apaturinae

*Dataset* – The dataset for the Apaturinae consisted of 9 taxa to which two outgroups were added: *Timelaea albescens* (Nymphalidae) and *Biblis hyperia (*Nymphalidae). We concatenated 10 gene fragments (COI, CAD, EF-1α, GAPDH, ArgK, IDH, MDH, RpS2, RpS5, wingless).

*Partition Finder* – Partition Finder identified 7 subsets.

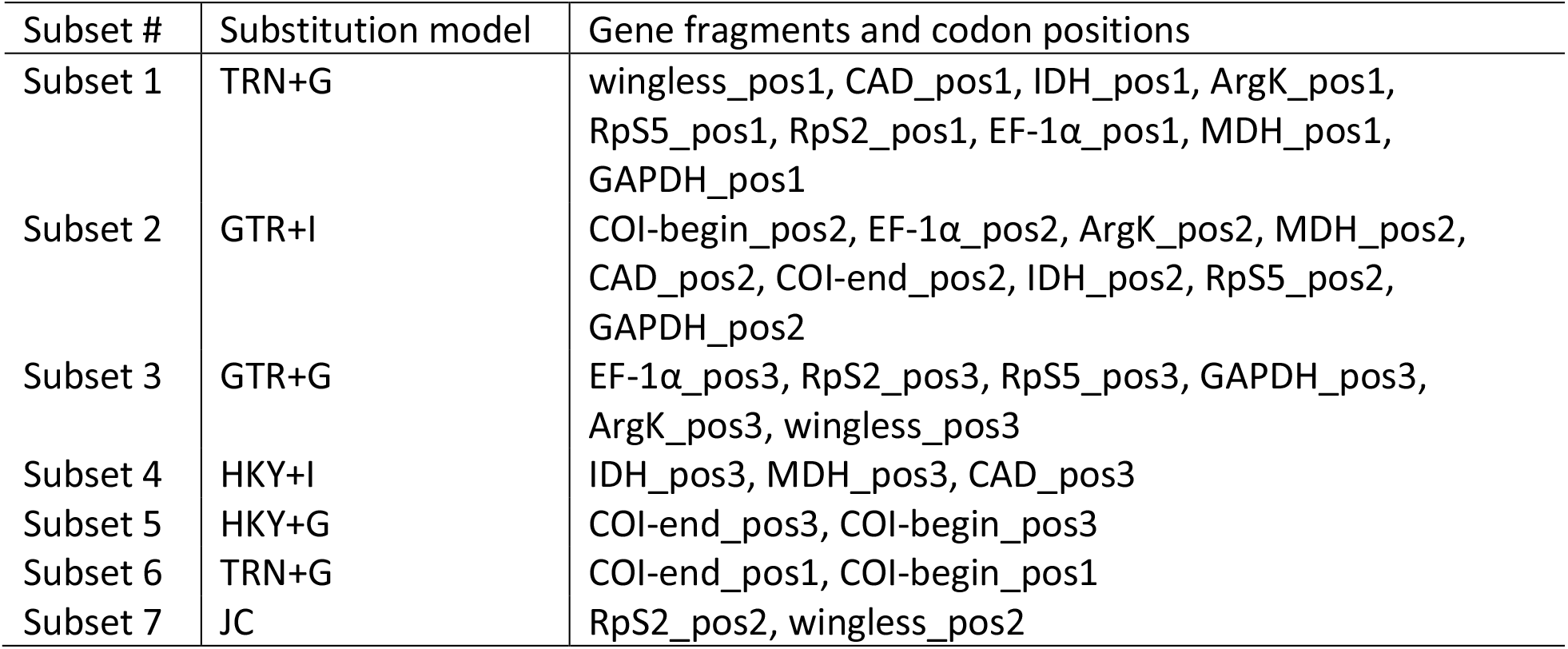

*BEAST analysis* – We used one molecular clock per subset identified by Partition Finder and obtained good mixing and convergence. We used a Birth-Death tree prior. We performed two runs of 20 million generations, sampling trees and parameters every 2000 generations.

*Grafting* – For grafting, the outgroups were removed and the subclade grafted at the mrca of Danainae.

#### 8- Nymphalidae: Heliconiinae + Limenitidinae

*Dataset –* The dataset combined the sister clades Heliconiinae and Limenitidinae and consisted of 92 taxa to which three outgroups were added: *Amnosia decora* (Nymphalidae), *Apatura iris* (Nymphalidae) and *Libythea celtis* (Nymphalidae). We concatenated 11 gene fragments (COI, CAD, EF-1α, GAPDH, ArgK, IDH, MDH, RpS2, RpS5, DDC, wingless).

*Partition Finder* – Partition Finder identified 14 subsets.

| Subset # | Substitution model | Gene fragments and codon positions |
| --- | --- | --- |
| Subset 1 | GTR+I+G | wingless_pos2, RpS2_pos1, ArgK_pos1 |
| Subset 2 | GTR+I+G | DDC_pos2, IDH_pos2, GAPDH_pos2, RpS5_pos2, EF-1α_pos2, ArgK_pos2 |
| Subset 3 | GTR+G | ArgK_pos3 |
| Subset 4 | HKY+I+G | MDH_pos3, CAD_pos3 |
| Subset 5 | GTR+I+G | DDC_pos1, wingless_pos1, CAD_pos1, RpS5_pos1, MDH_pos1, IDH_pos1 |
| Subset 6 | GTR+I+G | CAD_pos2, MDH_pos2, COI-begin_pos2, COI-end_pos2 |
| Subset 7 | GTR+I+G | COI-end_pos3, COI-begin_pos3 |
| Subset 8 | GTR+I+G | COI-end_pos1, COI-begin_pos1 |
| Subset 9 | GTR+I+G | DDC_pos3, wingless_pos3, RpS5_pos3, RpS2_pos3 |
| Subset 10 | GTR+I+G | EF-1α_pos3 |
| Subset 11 | TRN+I+G | EF-1α_pos1, GAPDH_pos1 |
| Subset 12 | GTR+I+G | GAPDH_pos3 |
| Subset 13 | GTR+I+G | IDH_pos3 |
| Subset 14 | JC+I | RpS2_pos2 |

*BEAST analysis* – Preliminary analyses showed problems with the subset 14 (RpS2_pos2) which was therefore removed from the analyses. In order to improve the quality of our runs we replaced the default priors for rates of substitutions by uniform priors ranging between 0 and 10 for the following case: subset7.cg. We used one molecular clock per subset identified by Partition Finder and obtained good mixing and convergence. We used a Birth-Death tree prior. We performed two runs of 100 million generations, sampling trees and parameters every 10000 generations.

*Grafting* – For grafting, the outgroups were removed and the subclade grafted at the split between Limenitidinae and Heliconiinae.

#### 9- Nymphalidae: Nymphalinae

*Dataset* – The dataset of Nymphalinae consisted of 83 taxa to which two outgroups were added: *Historis odius* (Nymphalidae) and *Pycina zamba* (Nymphalidae). We concatenated 11 gene fragments (COI, CAD, EF-1α, GAPDH, ArgK, IDH, MDH, RpS2, RpS5, DDC, wingless).

*Partition Finder* – Partition Finder identified 12 subsets.

| Subset # | Substitution model | Gene fragments and codon positions |
| --- | --- | --- |
| Subset 1 | TRN+I+G | wingless_pos1, ArgK_pos1, MDH_pos1, CAD_pos1, IDH_pos1, GAPDH_pos1 |
| Subset 2 | GTR+I+G | DDC_pos2, IDH_pos2, ArgK_pos2, EF-1α_pos2, RpS5_pos2, MDH_pos2, CAD_pos2 |
| Subset 3 | GTR+G | EF-1α_pos3, ArgK_pos3 |
| Subset 4 | HKY+I+G | MDH_pos3, CAD_pos3 |
| Subset 5 | GTR+G | COI-end_pos3, COI-begin_pos3 |
| Subset 6 | GTR+G | COI-end_pos1, COI-begin_pos1 |
| Subset 7 | GTR+I+G | COI-begin_pos2, COI-end_pos2 |
| Subset 8 | TRNEF | wingless_pos3, DDC_pos3 |
| Subset 9 | SYM+G | DDC_pos1, wingless_pos2 |
| Subset 10 | GTR+I+G | RpS5_pos1, RpS2_pos1, EF-1α_pos1 |
| Subset 11 | GTR+G | IDH_pos3, RpS2_pos3, GAPDH_pos3, RpS5_pos3 |
| Subset 12 | JC+I | RpS2_pos2, GAPDH_pos2 |

*BEAST analysis* – In order to improve the quality of our runs we replaced the default priors for rates of substitutions by uniform priors ranging between 0 and 10 for the following case: subset5.cg. Preliminary analyses revealed problems when using one molecular clock per subset identified by Partition Finder. We restricted the analysis to one molecular clock for the mitochondrial gene fragments and one molecular clock for the nuclear gene fragments. We used a Birth-Death tree prior. We performed two runs of 100 million generations, sampling trees and parameters every 10000 generations.

*Grafting* – For grafting, the outgroups were removed and the subclade grafted at the mrca of Nymphalinae.

#### 10- Nymphalidae: Satyrinae 1

*Dataset* – The first Satyrinae dataset consisted of 13 taxa, belonging to the genera *Kirinia*, *Pararge*, *Lasiommata*, *Tatinga*, *Chonala* and *Lopinga*, to which three outgroups were added: *Bicyclus anynana* (Nymphalidae), *Acrophtalmia leuce* (Nymphalidae) and *Ragadia makuta* (Nymphalidae). We concatenated 5 gene fragments (COI, EF-1α, GAPDH, RpS5, wingless).

*Partition Finder* – Partition Finder identified 6 subsets.

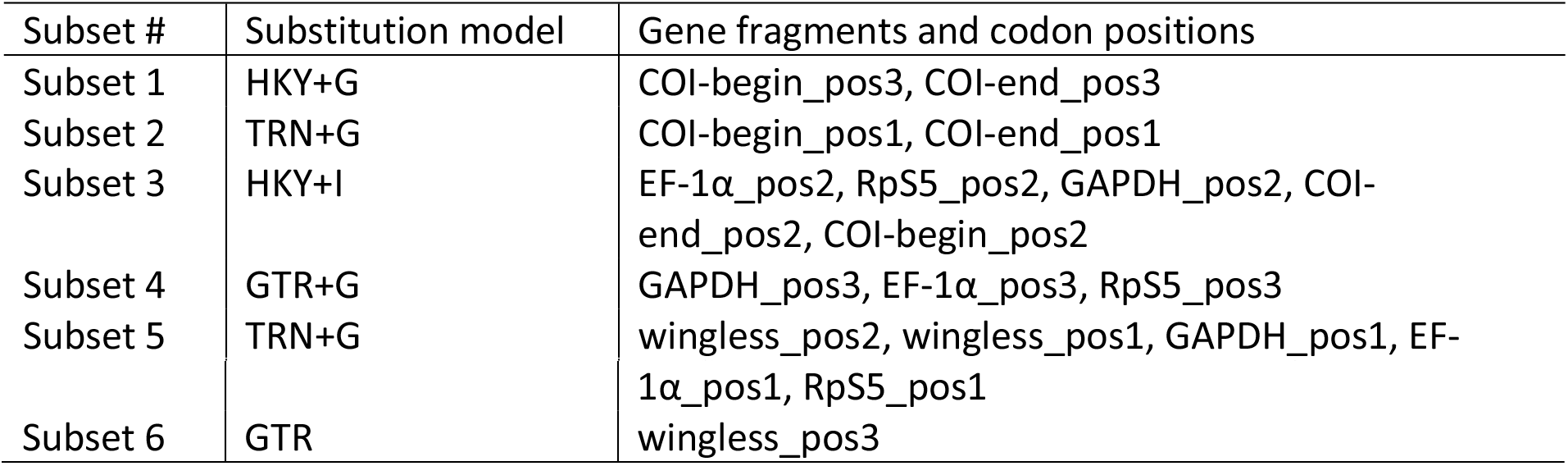

*BEAST analysis* – We used one molecular clock per subset identified by Partition Finder and obtained good mixing and convergence. We used a Birth-Death tree prior. We performed two runs of 20 million generations, sampling trees and parameters every 2000 generations.

*Grafting* – For grafting, the outgroups were removed and the subclade grafted at the crown of the clade after removing the outgroups.

#### 11- Nymphalidae: Satyrinae 2

*Dataset* – The second Satyrinae dataset consisted of 161 taxa, belonging to the genera *Calisto, Euptychia, Callerebia, Proterebia, Gyrocheilus, Strabena, Ypthima, Ypthimomorpha*, *Stygionympha*, *Cassionympha*, *Neocoenyra*, *Pseudonympha*, *Erebia*, *Boerebia*, *Hyponephele*, *Cercyonis*, *Maniola*, *Aphantopus*, *Pyronia*, *Faunula*, *Grumia*, *Paralasa*, *Melanargia*, *Hipparchia*, *Berberia*, *Oeneis*, *Neominois*, *Karanasa*, *Brintesia*, *Arethusana*, *Satyrus*, *Pseudochazara* and *Chazara*, to which three outgroups were added: *Coenonympha pamphilus* (Nymphalidae), *Taygetis virgilia* (Nymphalidae) and *Pronophila thelebe* (Nymphalidae). We concatenated 10 gene fragments (COI, CAD, EF-1α, GAPDH, ArgK, IDH, MDH, RpS2, RpS5, wingless).

*Partition Finder* – Partition Finder identified 11 subsets.

| Subset # | Substitution model | Gene fragments and codon positions |
| --- | --- | --- |
| Subset 1 | GTR+I+G | ArgK_pos1, wingless_pos1, RpS2_pos1, IDH_pos1, CAD_pos1, EF-1α_pos1, GAPDH_pos1, RpS5_pos1, MDH_pos1 |
| Subset 2 | HKY+I+G | MDH_pos2, ArgK_pos2, CAD_pos2, IDH_pos2, RpS5_pos2, EF-1α_pos2, GAPDH_pos2 |
| Subset 3 | GTR+G | ArgK_pos3, wingless_pos3 |
| Subset 4 | HKY+G | IDH_pos3, MDH_pos3, CAD_pos3 |
| Subset 5 | GTR+G | COI-begin_pos3, COI-end_pos3 |
| Subset 6 | GTR+G | COI-begin_pos1, COI-end_pos1 |
| Subset 7 | TRN+I+G | COI-begin_pos2, COI-end_pos2 |
| Subset 8 | GTR+I+G | EF-1α_pos3, RpS5_pos3 |
| Subset 9 | HKY+G | GAPDH_pos3 |
| Subset 10 | TRNEF+G | RpS2_pos2, wingless_pos2 |
| Subset 11 | K80+G | RpS2_pos3 |

*BEAST analysis* – In order to improve the quality of our runs we replaced the default priors for rates of substitutions by uniform prior ranging between 0 and 10 for the following cases: subset5.ac, subset5.ag, subset5.at, subset5.cg, subset5.gt. We used one molecular clock per subset identified by Partition Finder and obtained good mixing and convergence. We used a Birth-Death tree prior. We performed two runs of 100 million generations, sampling trees and parameters every 10000 generations.

*Grafting* – For grafting, the outgroups were removed and the subclade grafted at the crown of the clade after removing the outgroups.

#### 12- Nymphalidae: Satyrinae 3

*Dataset* – The third Satyrinae dataset consisted of 15 taxa all belonging to the genus *Coenonympha*, to which two outgroups were added: *Sinonympha amoena* (Nymphalidae) and *Oressinoma sorata* (Nymphalidae). We concatenated 9 gene fragments (COI, CAD, EF-1α, GAPDH, IDH, MDH, RpS2, RpS5, wingless).

*Partition Finder* – Partition Finder identified 6 subsets.

| Subset # | Substitution model | Gene fragments and codon positions |
| --- | --- | --- |
| Subset 1 | HKY | CAD_pos3 |
| Subset 2 | TRN+G | RpS2_pos2, COI-begin_pos1, COI-end_pos1, CAD_pos1, IDH_pos1 |
| Subset 3 | HKY | RpS5_pos2, MDH_pos2, GAPDH_pos2, EF-1α_pos2, IDH_pos2, CAD_pos2 |
| Subset 4 | GTR+G | COI-end_pos3, COI-begin_pos3 |
| Subset 5 | HKY+I | COI-end_pos2, COI-begin_pos2 |
| Subset 6 | HKY+G | wingless_pos3, RpS2_pos3, EF-1α_pos3 |
| Subset 7 | TRN+G | wingless_pos1, RpS2_pos1, MDH_pos1, GAPDH_pos1, RpS5_pos1, EF-1α_pos1 |
| Subset 8 | HKY+G | RpS5_pos3, MDH_pos3, GAPDH_pos3, IDH_pos3 |
| Subset 9 | JC | wingless_pos2 |

*BEAST analysis* – We used one molecular clock per subset identified by Partition Finder and obtained good mixing and convergence. We used a Birth-Death tree prior. We performed two runs of 20 million generations, sampling trees and parameters every 2000 generations.

*Grafting* – For grafting, the outgroups were removed and the subclade grafted at the crown of *Coenonympha*.

## Data Records

The analysed dataset (a concatenated alignment of the genes COI, CAD, EF-1α, GAPDH, ArgK, IDH, MDH, RpS2, RpS5, DDC, wingless, and H3) is available in FASTA format at DOI: 10.5281/zenodo.3531555. The posterior distribution of ML trees and the consensus tree are available in NEWICK format at DOI: 10.5281/zenodo.3531555.

## Technical Validation

Species identities of the chosen sequences for the dataset were validated by blasting the DNA barcode sequence against the Barcode Of Life Database, which has a good representation of European butterfly species due to a number of barcoding projects implemented in different countries. In almost all cases, the sequences came from the same voucher specimen itself. In 88 cases (Supporting Information), the sequences used were from different individuals. In these cases special care was taken to use sequences from reliable sources, preferably those with voucher photographs.

We based our time-calibration from a recent re-evaluation of the timing of divergence of higher-level Papilionoidea. We used the topology inferred by Chazot et al. ^47^ as a backbone in our grafting procedure. This topology was fixed in Chazot et al. ^47^, hence only node ages were estimated. Within each subclade we grafted however, we let BEAST estimate the topology in addition to node height. Several sections of the European butterfly tree remain poorly supported. This most likely arises from the lack of molecular information as well as recent and rapid diversification events within *Polyommatus*, *Hipparchia*, or *Pseudochazara* for example. We show here a synthetic tree summarizing the posterior distribution of topologies and node ages but the posterior distribution of grafted trees can be found in Supporting Information, providing a distribution of alternative topologies and node ages estimated by BEAST. We strongly advise any researcher using our phylogenetic framework to repeat the analyses on at least 100 trees randomly sampled from this posterior distribution in order to account for topology and node age uncertainty. This tree can also help identify the sections of the tree lacking molecular information and therefore points at the sections that should be targeted in the future when generating new molecular data.

## Usage Notes

We have generated a robust phylogenetic hypothesis for all European species of butterflies with associated times of divergence (Fig. 1). Our purpose is to provide a complete phylogenetic framework for use by the ecological and evolutionary communities. The demand for such a phylogenetic information is high at the moment and various proxies have been used that are not ideal, starting already in 2005^88^. We provide a posterior distribution of topologies and node ages, in order for researchers to be able to take phylogenetic and node age uncertainty into account if they so wish. The tree files are provided in standard Newick format as output from BEAST. Future studies will not necessarily be as comprehensive as the tree we provide. In such cases we recommend using tools such as the *ape* package^89^ in R^90^ to remove tips from the tree that are not relevant to a given study.

## Acknowledgements

MW thanks Brigitte Gottsberger (University of Vienna) for assistance in the lab and the following colleagues for specimen samples or sequences: Benedicto Acosta-Fernandez (Spain), Bernard Turlin (France), Dirk Gerber (Germany), Eddie John (UK), John Coutsis (Greece), Javier García (Spain), Karen van Dorp (Netherlands), Klaus Schurian (Germany), Pedro Oromí (Spain), Peter Russell (UK), Roger Vila (Spain), Vlad Dinca (Finland), Xavier Merit (France), Zdenek Fric (Czech Republic), and Zdravko Kolev (Bulgaria). NW acknowledges funding from the Department of Biology, Lund University, the Swedish Research Council (grant number 2015-04441). NC acknowledges funding from BECC (Biodiversity and Ecosystem services in a Changing Climate). CWW acknowledges funding from the Swedish Research Council (grant number 2017-04386). The study was also supported by iDiv through the sDiv working group sECURE (https://www.idiv.de/secure).

---

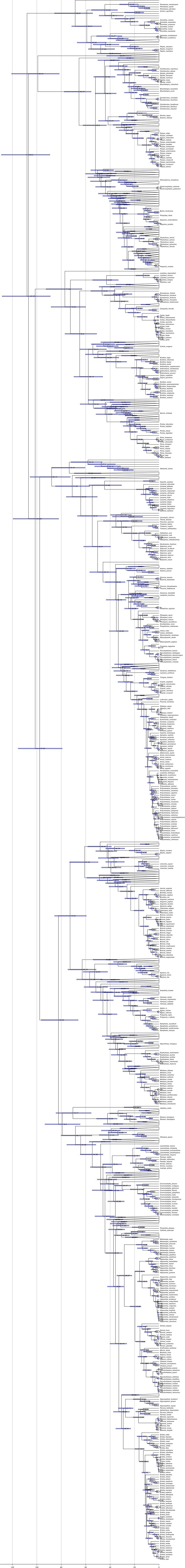

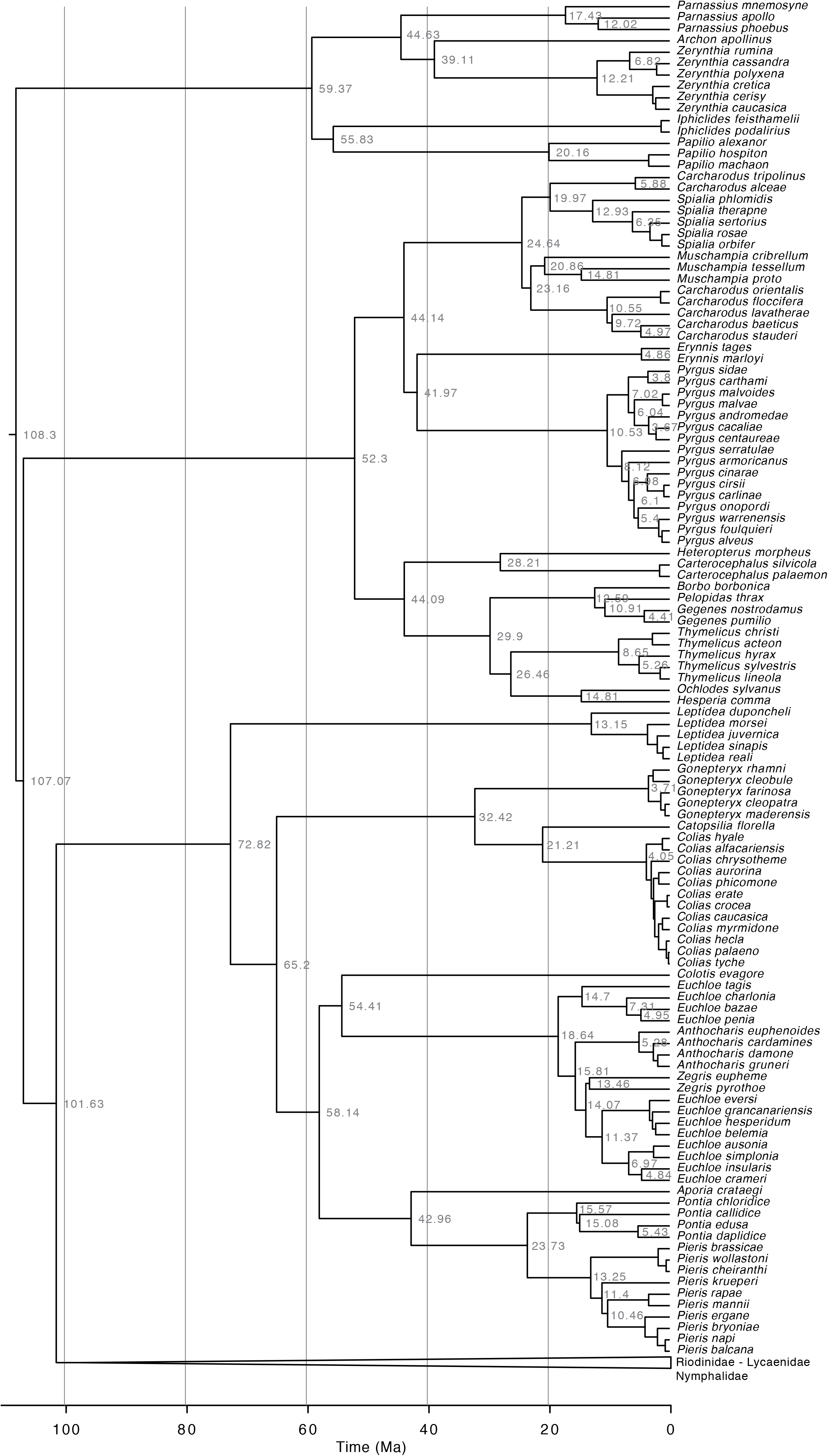

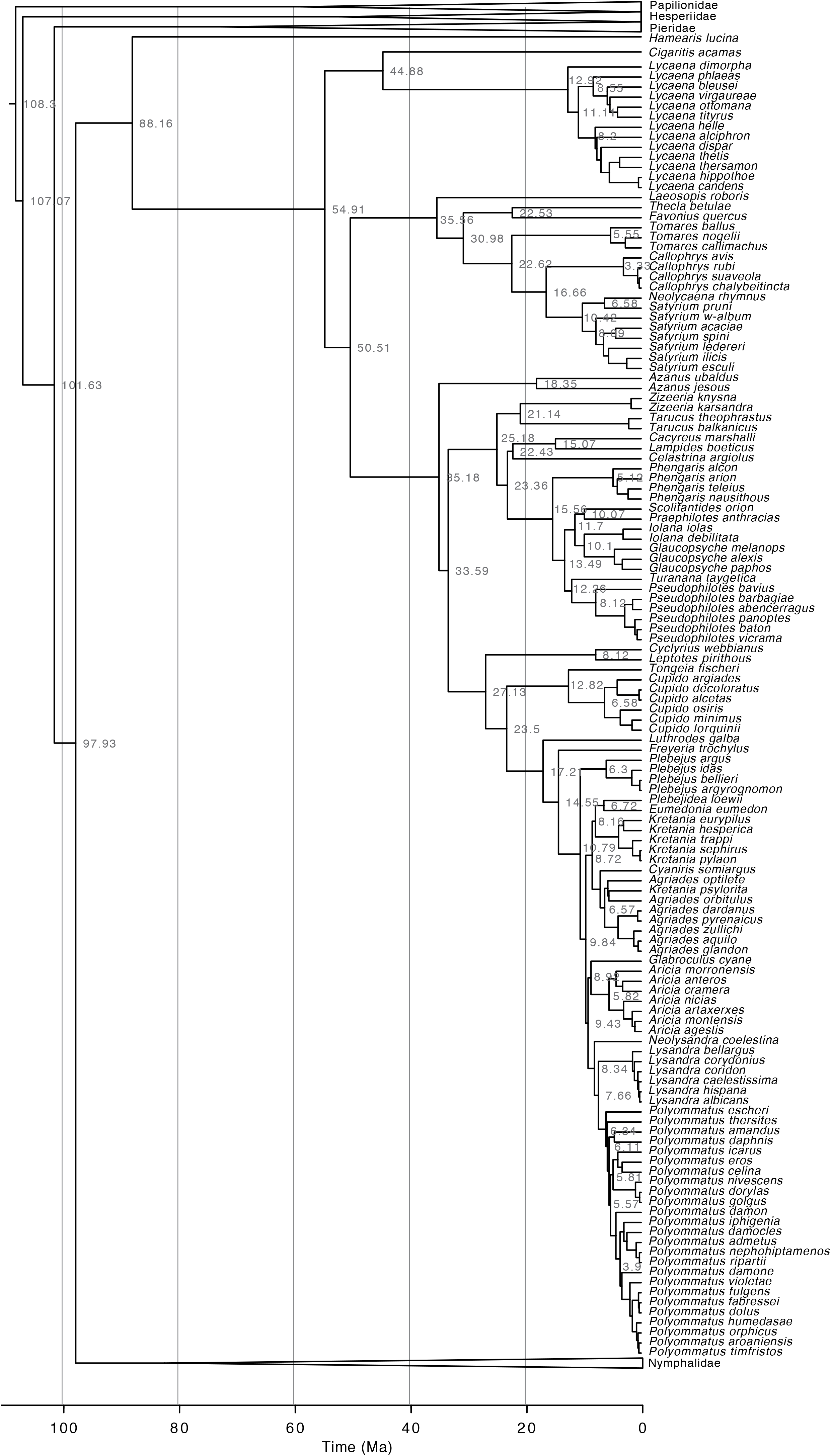

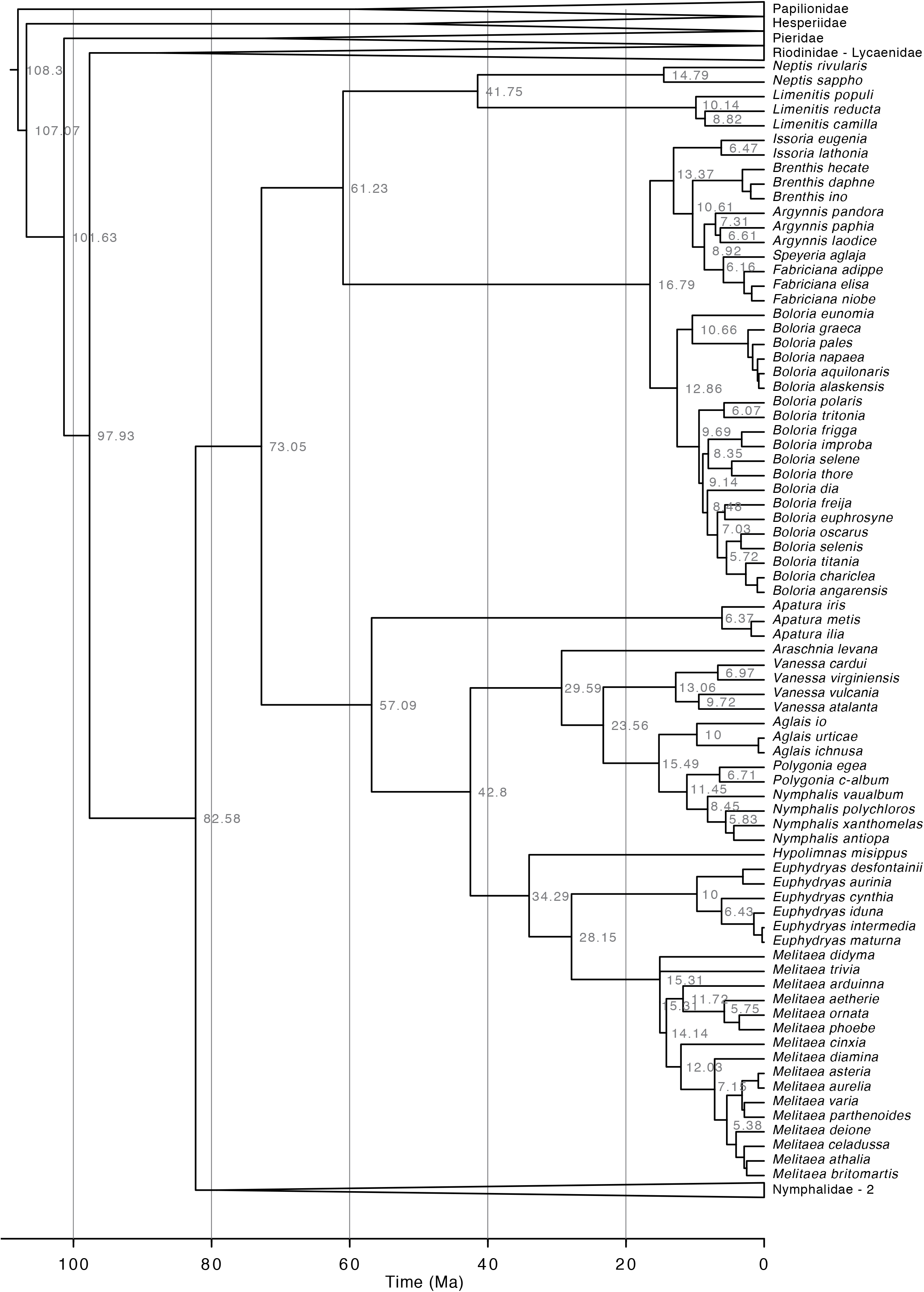

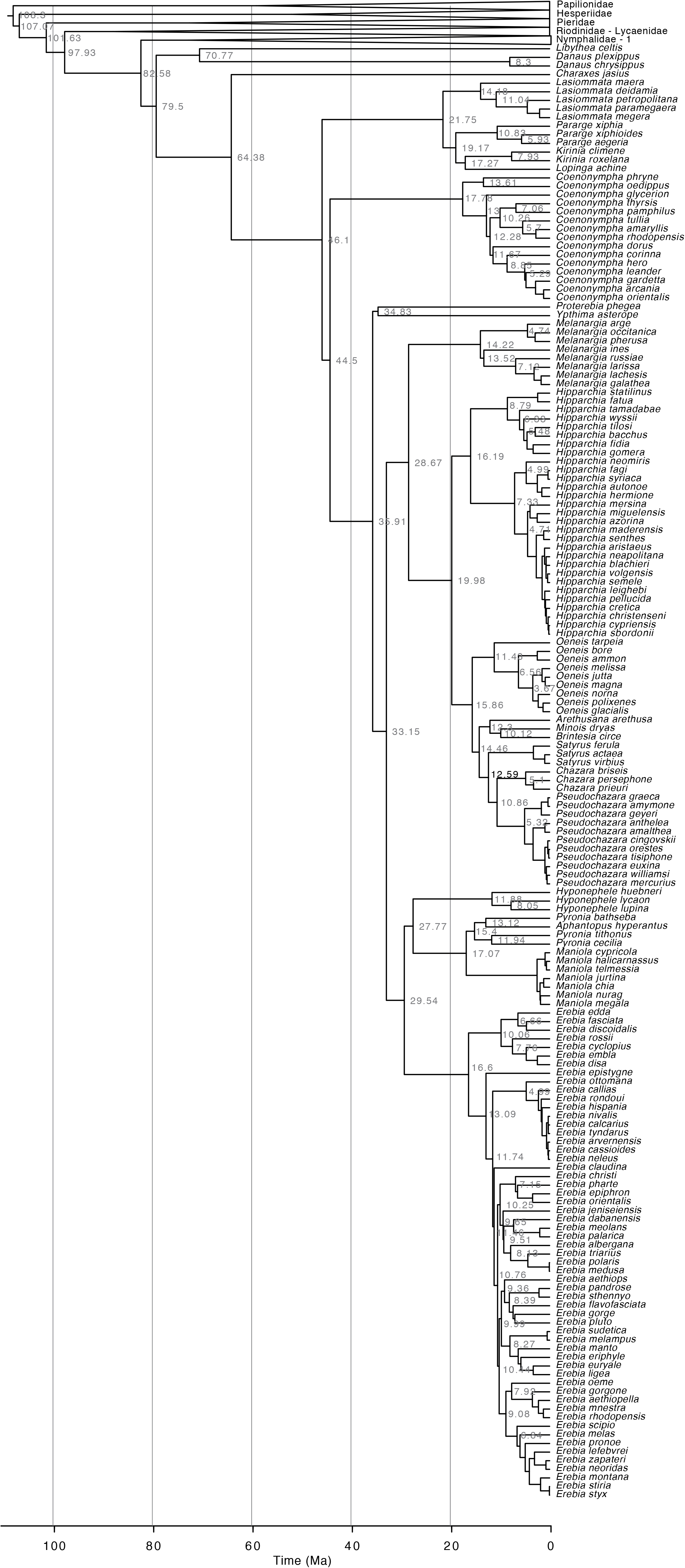

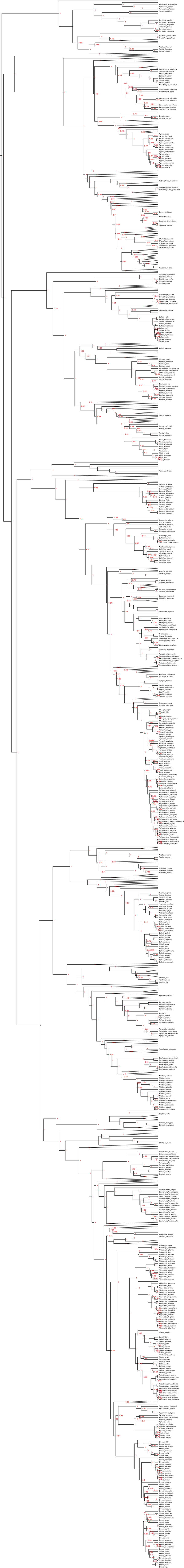

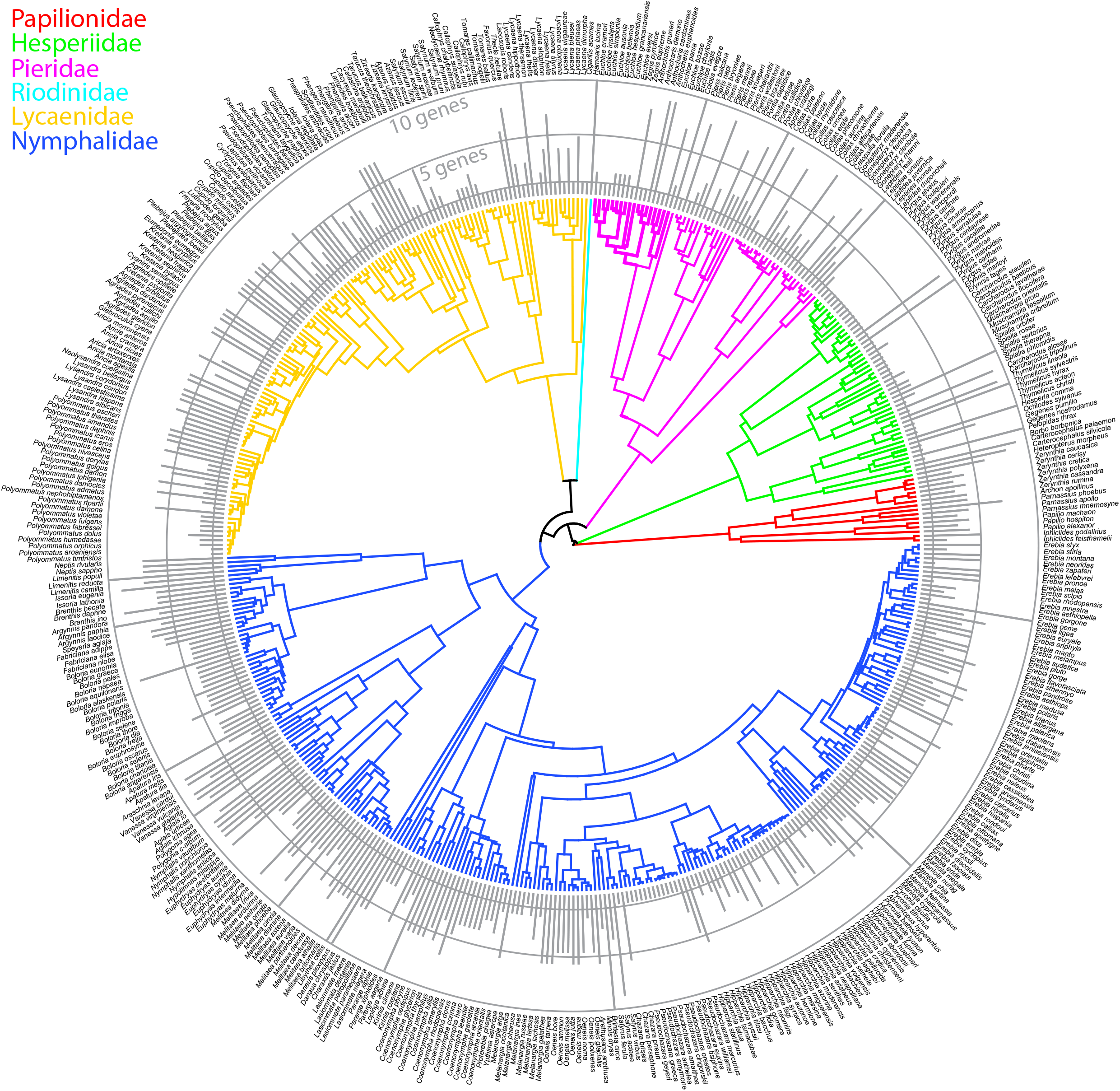

## References

1 Mouquet, N. et al. Ecophylogenetics: advances and perspectives. Biol Rev 87, 769–785 doi:10.1111/j.1469-185X.2012.00224.x (2012).

2 Webb, C. O., Ackerly, D. D., McPeek, M. A. & Donoghue, M. J. Phylogenies and community ecology. Annu Rev Ecol Syst 33, 475–505 doi:10.1146/annurev.ecolysis.33.010802.150448 (2002).

3 Roquet, C., Thuiller, W. & Lavergne, S. Building megaphylogenies for macroecology: taking up the challenge. Ecography 36, 13–26 doi:10.1111/j.1600-0587.2012.07773.x (2013).

4 De Palma, A. et al. Dimensions of biodiversity loss: Spatial mismatch in land-use impacts on species, functional and phylogenetic diversity of European bees. Divers Distrib 23, 1435–1446 doi:doi:10.1111/ddi.12638 (2017).

5 Mazel, F. et al. Global patterns of β-diversity along the phylogenetic time-scale: The role of climate and plate tectonics. Global Ecol Biogeogr 26, 1211–1221 doi:doi:10.1111/geb.12632 (2017).

6 Economo, E. P., Narula, N., Friedman, N. R., Weiser, M. D. & Guénard, B. Macroecology and macroevolution of the latitudinal diversity gradient in ants. Nature Communications 9, 1778, doi:10.1038/s41467-018-04218-4 (2018).

7 Mazel, F. et al. Mammalian phylogenetic diversity-area relationships at a continental scale. Ecology 96, 2814–2822 (2015).

8 Cavender-Bares, J., Kozak, K. H., Fine, P. V. A. & Kembel, S. W. The merging of community ecology and phylogenetic biology. Ecol Lett 12, 693–715 doi:doi:10.1111/j.1461-0248.2009.01314.x (2009).

9 D’Amen, M. et al. Improving spatial predictions of taxonomic, functional and phylogenetic diversity. J Ecol 106, 76–86, doi:doi:10.1111/1365-2745.12801 (2018).

10 Gerhold, P., Cahill, J. F., Winter, M., Bartish, I. V. & Prinzing, A. Phylogenetic patterns are not proxies of community assembly mechanisms (they are far better). Funct Ecol 29, 600–614 doi:10.1111/1365-2435.12425 (2015).

11 Monnet, A. C. et al. Asynchrony of taxonomic, functional and phylogenetic diversity in birds. Global Ecol Biogeogr 23, 780–788 doi:10.1111/geb.12179 (2014).

12 Ovaskainen, O. et al. How to make more out of community data? A conceptual framework and its implementation as models and software. Ecol Lett, n/a-n/a, doi:10.1111/ele.12757 (2017).

13 Gallien, L., Altermatt, F., Wiemers, M., Schweiger, O. & Zimmermann, N. E. Invasive plants threaten the least mobile butterflies in Switzerland. Divers Distrib 23, 185–195 doi:10.1111/ddi.12513 (2017).

14 Winter, M. et al. Plant extinctions and introductions lead to phylogenetic and taxonomic homogenization of the European flora. P Natl Acad Sci USA 106, 21721–21725 (2009).

15 Li, S.-p. et al. The effects of phylogenetic relatedness on invasion success and impact: deconstructing Darwin’s naturalisation conundrum. Ecol Lett 18, 1285–1292 doi:doi:10.1111/ele.12522 (2015).

16 Knapp, S., Kühn, I., Schweiger, O. & Klotz, S. Challenging urban species diversity: contrasting phylogenetic patterns across plant functional groups in Germany. Ecol Lett 11, 1054–1064 (2008).

17 Díaz, S. et al. Functional traits, the phylogeny of function, and ecosystem service vulnerability. Ecol Evol 3, 2958–2975, doi:doi:10.1002/ece3.601 (2013).

18 Davies, T. J., Urban, M. C., Rayfield, B., Cadotte, M. W. & Peres-Neto, P. R. Deconstructing the relationships between phylogenetic diversity and ecology: a case study on ecosystem functioning. Ecology 97, 2212–2222 doi:doi:10.1002/ecy.1507 (2016).

19 Mazel, F. et al. Prioritizing phylogenetic diversity captures functional diversity unreliably. Nature Communications 9, doi:10.1038/s41467-018-05126-3 (2018).

20 Winter, M., Devictor, V. & Schweiger, O. Phylogenetic diversity and nature conservation: where are we? Trends Ecol Evol 28, 199–204 doi:10.1016/j.tree.2012.10.015 (2013).

21 Wiens, J. J. & Graham, C. H. Niche Conservatism: Integrating Evolution, Ecology, and Conservation Biology. Annual Review of Ecology, Evolution, and Systematics 36, 519–539 doi:10.1146/annurev.ecolsys.36.102803.095431 (2005).

22 Kühn, I., Nobis, M. P. & Durka, W. Combining spatial and phylogenetic eigenvector filtering in trait analysis. Global Ecol Biogeogr 18, 745–758 (2009).

23 Schweiger, O., Klotz, S., Durka, W. & Kühn, I. A comparative test of phylogenetic diversity indices. Oecologia 157, 485–495 doi:10.1007/s00442-008-1082-2 (2008).

24 Tucker, C. M. et al. A guide to phylogenetic metrics for conservation, community ecology and macroecology. Biol Rev, n/a-n/a, doi:10.1111/brv.12252 (2016).

25 Morales-Castilla, I., Davies, T. J., Pearse, W. D. & Peres-Neto, P. Combining phylogeny and co-occurrence to improve single species distribution models. Global Ecol Biogeogr 26, 740–752 doi:doi:10.1111/geb.12580 (2017).

26 Lavergne, S., Evans, M. E. K., Burfield, I. J., Jiguet, F. & Thuiller, W. Are species’ responses to global change predicted by past niche evolution? Philos T R Soc B 368, doi:10.1098/rstb.2012.0091 (2013).

27 Thuiller, W. et al. Consequences of climate change on the tree of life in Europe. Nature 470, 531–534 doi:10.1038/nature09705 (2011).

28 Thuiller, W. et al. Conserving the functional and phylogenetic trees of life of European tetrapods. Philosophical Transactions of the Royal Society B: Biological Sciences 370, 20140005, doi:10.1098/rstb.2014.0005 (2015).

29 Durka, W. & Michalski, S. G. Daphne: a dated phylogeny of a large European flora for phylogenetically informed ecological analyses. Ecology 93, 2297 (2012).

30 Jetz, W., Thomas, G. H., Joy, J. B., Hartmann, K. & Mooers, A. O. The global diversity of birds in space and time. Nature 491, 444–448 doi:10.1038/nature11631 (2012).

31 Roquet, C., Lavergne, S. & Thuiller, W. One tree to link them all: a phylogenetic dataset for the European tetrapoda. PLoS Curr 6, doi:10.1371/currents.tol.5102670fff8aa5c918e78f5592790e48 (2014).

32 Stork, N. E. How Many Species of Insects and Other Terrestrial Arthropods Are There on Earth? Annual Review of Entomology, Vol 63 63, 31–45 doi:10.1146/annurev-ento-020117-043348 (2018).

33 Noriega, J. A. et al. Research trends in ecosystem services provided by insects. Basic Appl Ecol 26, 8–23 doi:10.1016/j.baae.2017.09.006 (2018).

34 McGeoch, M. A. in Conservation Biology (eds A. J. A. Stewart, T. R. New, & O. T. Lewis) 144–174 (CABI Publishing, 2007).

35 Wiemers, M. et al. An updated checklist of the European Butterflies (Lepidoptera, Papilionoidea). Zookeys, 9–45, doi:10.3897/zookeys.811.28712 (2018).

36 Settele, J., Shreeve, T. G., Konvicka, M. & van Dyck, H. in Ecology of butterflies in Europe xii + 513 (Cambridge University Press, Cambridge, 2009).

37 Devictor, V. et al. Differences in the climatic debts of birds and butterflies at a continental scale. Nature Climate Change 2, 121–124 (2012).

38 Schweiger, O., Harpke, A., Wiemers, M. & Settele, J. CLIMBER: Climatic niche characteristics of the butterflies in Europe. ZooKeys 367, 65–84 doi:DOI 10.3897/zookeys.367.6185 (2014).

39 Bartonova, A., Benes, J. & Konvicka, M. Generalist-specialist continuum and life history traits of Central European butterflies (Lepidoptera) - are we missing a part of the picture? Eur J Entomol 111, 543–553 doi:10.14411/eje.2014.060 (2014).

40 van Swaay, C. et al. European Red List of Butterflies. (Publications Office of the European Union, 2010).

41 van Swaay, C., Warren, M. & Lois, G. Biotope use and trends of European butterflies. J Insect Conserv 10, 305–306 doi:10.1007/s10841-006-8361-1 (2006).

42 Settele, J. et al. Climatic risk atlas of European butterflies. BioRisk 1, 1–710 (2008).

43 Kudrna, O. et al. Distribution Atlas of Butterflies in Europe. (Gesellschaft für Schmetterlingsschutz, 2011).

44 Espeland, M. et al. A Comprehensive and Dated Phylogenomic Analysis of Butterflies. Curr Biol 28, 770-+, doi:10.1016/j.cub.2018.01.061 (2018).

45 Heikkila, M., Kaila, L., Mutanen, M., Pena, C. & Wahlberg, N. Cretaceous origin and repeated tertiary diversification of the redefined butterflies. P R Soc B 279, 1093–1099 doi:10.1098/rspb.2011.1430 (2012).

46 Wahlberg, N. et al. Synergistic effects of combining morphological and molecular data in resolving the phylogeny of butterflies and skippers. P R Soc B 272, 1577–1586 doi:10.1098/rspb.2005.3124 (2005).

47 Chazot, N. et al. Priors and Posteriors in Bayesian Timing of Divergence Analyses: the Age of Butterflies Revisited. Syst Biol 68, 797–813 doi:10.1093/sysbio/syz002 (2019).

48 Braby, M. F., Vila, R. & Pierce, N. E. Molecular phylogeny and systematics of the Pieridae (Lepidoptera: Papilionoidea): higher classification and biogeography. Zoological Journal of the Linnean Society 147, 239–275 doi:10.1111/j.1096-3642.2006.00264.x (2006).

49 Campbell, D. L., Brower, A. V. & Pierce, N. E. Molecular evolution of the wingless gene and its implications for the phylogenetic placement of the butterfly family Riodinidae (Lepidoptera: Papilionoidea). Mol Biol Evol 17, 684–696 doi:10.1093/oxfordjournals.molbev.a026347 (2000).

50 Caterino, M. S., Reed, R. D., Kuo, M. M. & Sperling, F. A. H. A Partitioned Likelihood Analysis of Swallowtail Phylogeny (Lepidoptera: Papilionidae). Syst. Biol. 50, 106–127 doi:10.1080/106351501750107530 (2001).

51 Wahlberg, N., Weingartner, E. & Nylin, S. Towards a better understanding of the higher systematics of Nymphalidae (Lepidoptera: Papilionoidea). Mol Phylogenet Evol 28, 473–484 doi:10.1016/S1055-7903(03)00052-6 (2003).

52 Warren, A. D., Ogawa, J. R. & Brower, A. V. Z. Phylogenetic relationships of subfamilies and circumscription of tribes in the family Hesperiidae (Lepidoptera: Hesperioidea). Cladistics 24, 642–676 doi:10.1111/j.1096-0031.2008.00218.x (2008).

53 Espeland, M. et al. Ancient Neotropical origin and recent recolonisation: Phylogeny, biogeography and diversification of the Riodinidae (Lepidoptera: Papilionoidea). Mol Phylogenet Evol 93, 296–306 doi:10.1016/j.ympev.2015.08.006 (2015).

54 Sahoo, R. K. et al. Ten genes and two topologies: an exploration of higher relationships in skipper butterflies (Hesperiidae). Peerj 4, doi:10.7717/peerj.2653 (2016).

55 Seraphim, N. et al. Molecular phylogeny and higher systematics of the metalmark butterflies (Lepidoptera: Riodinidae). Syst Entomol 43, 407–425 doi:10.1111/syen.12282 (2018).

56 Wahlberg, N. et al. Nymphalid butterflies diversify following near demise at the Cretaceous/Tertiary boundary. P R Soc B 276, 4295–4302 doi:10.1098/rspb.2009.1303 (2009).

57 Wahlberg, N., Rota, J., Braby, M. F., Pierce, N. E. & Wheat, C. W. Revised systematics and higher classification of pierid butterflies (Lepidoptera: Pieridae) based on molecular data. Zool Scr 43, 641–650 doi:10.1111/zsc.12075 (2014).

58 Allio, R. et al. Whole genome shotgun phylogenomics resolves the pattern and timing of swallowtail butterfly evolution. Syst Biol, doi:10.1093/sysbio/syz030 (2019).

59 Nylin, S. & Wahlberg, N. Does plasticity drive speciation? Host-plant shifts and diversification in nymphaline butterflies (Lepidoptera: Nymphalidae) during the tertiary. Biol. J. Linn. Soc. 94, 115–130 doi:10.1111/j.1095-8312.2008.00964.x (2008).

60 Pena, C. & Wahlberg, N. Prehistorical climate change increased diversification of a group of butterflies. Biol Letters 4, 274–278 doi:10.1098/rsbl.2008.0062 (2008).

61 Pena, C. et al. Higher level phylogeny of Satyrinae butterflies (Lepidoptera: Nymphalidae) based on DNA sequence data. Mol Phylogenet Evol 40, 29–49 doi:10.1016/j.ympev.2006.02.007 (2006).

62 Wahlberg, N., Brower, A. V. Z. & Nylin, S. Phylogenetic relationships and historical biogeography of tribes and genera in the subfamily Nymphalinae (Lepidoptera: Nymphalidae). Biol. J. Linn. Soc. 86, 227–251 doi:DOI 10.1111/j.1095-8312.2005.00531.x (2005).

63 Talavera, G., Lukhtanov, V. A., Pierce, N. E. & Vila, R. Establishing criteria for higher-level classification using molecular data: the systematics of Polyommatus blue butterflies (Lepidoptera, Lycaenidae). Cladistics 29, 166–192, doi:10.1111/j.1096-0031.2012.00421.x (2013).

64 Wiemers, M., Stradomsky, B. V. & Vodolazhsky, D. I. A molecular phylogeny of *Polyommatus* s. str. and *Plebicula* based on mitochondrial COI and nuclear ITS2 sequences (Lepidoptera: Lycaenidae). Eur J Entomol 107, 325–336 (2010).

65 Pena, C., Witthauer, H., Kleckova, I., Fric, Z. & Wahlberg, N. Adaptive radiations in butterflies: evolutionary history of the genus Erebia (Nymphalidae: Satyrinae). Biol. J. Linn. Soc. 116, 449–467 doi:10.1111/bij.12597 (2015).

66 Wiemers, M. & Fiedler, K. Does the DNA barcoding gap exist? - a case study in blue butterflies (Lepidoptera: Lycaenidae). Front Zool 4, 8, doi:10.1186/1742-9994-4-8 (2007).

67 Dinca, V. et al. DNA barcode reference library for Iberian butterflies enables a continental-scale preview of potential cryptic diversity. Sci Rep-Uk 5, doi:10.1038/srep12395 (2015).

68 Dincă, V., Zakharov, E., Hebert, P. D. & Vila, R. Complete DNA barcode reference library for a country’s butterfly fauna reveals high performance for temperate Europe. P R Soc B 278, 347–355 doi:10.1098/rspb.2010.1089 (2011).

69 Hausmann, A. et al. Now DNA-barcoded: the butterflies and larger moths of Germany. Spixiana 34, 47–58 (2011).

70 Huemer, P. & Wiesmair, B. in Wissenschaftliches Jahrbuch der Tiroler Landesmuseen 2017 8–33 (StudienVerlag, 2017).

71 Litman, J. et al. A DNA barcode reference library for Swiss butterflies and forester moths as a tool for species identification, systematics and conservation. Plos One 13, doi:10.1371/journal.pone.0208639 (2018).

72 Bowler, D. E. et al. Cross-realm assessment of climate change impacts on species’ abundance trends. Nat Ecol Evol 1, doi:10.1038/s41559-016-0067 (2017).

73 Bowler, D. E. et al. A cross-taxon analysis of the impact of climate change on abundance trends in central Europe. Biol Conserv 187, 41–50 doi:10.1016/j.biocon.2015.03.034 (2015).

74 Schleuning, M. et al. Ecological networks are more sensitive to plant than to animal extinction under climate change. Nature Communications 7, doi:10.1038/ncomms13965 (2016).

75 Essens, T., van Langevelde, F., Vos, R. A., Van Swaay, C. A. M. & WallisDeVries, M. F. Ecological determinants of butterfly vulnerability across the European continent. J Insect Conserv 21, 439–450 doi:10.1007/s10841-017-9972-4 (2017).

76 Dapporto, L. et al. Integrating three comprehensive data sets shows that mitochondrial DNA variation is linked to species traits and paleogeographic events in European butterflies. Mol Ecol Resour 19, 1623–1636, doi:10.1111/1755-0998.13059 (2019).

77 Pena, C. & Malm, T. VoSeq: A Voucher and DNA Sequence Web Application. Plos One 7, e39071, doi:10.1371/journal.pone.0039071 (2012).

78 Wahlberg, N. & Wheat, C. W. Genomic outposts serve the phylogenomic pioneers: Designing novel nuclear markers for genomic DNA extractions of lepidoptera. Syst Biol 57, 231–242 doi:10.1080/10635150802033006 (2008).

79 Fric, Z. F. et al. World travellers: phylogeny and biogeography of the butterfly genus *Leptotes* (Lepidoptera: Lycaenidae). Syst Entomol 44, 652–665 doi:10.1111/syen.12349 (2019).

80 Kawahara, A. Y. et al. Phylogenomics reveals the evolutionary timing and pattern of butterflies and moths. Proc Natl Acad Sci U S A, doi:10.1073/pnas.1907847116 (2019).

81 Stamatakis, A. RAxML version 8: a tool for phylogenetic analysis and post-analysis of large phylogenies. Bioinformatics 30, 1312–1313 doi:10.1093/bioinformatics/btu033 (2014).

82 Suchard, M. A. et al. Bayesian phylogenetic and phylodynamic data integration using BEAST 1.10. Virus Evolution 4, vey016–vey016, doi:10.1093/ve/vey016 (2018).

83 Aduse-Poku, K., Vingerhoedt, E. & Wahlberg, N. Out-of-Africa again: a phylogenetic hypothesis of the genus Charaxes (Lepidoptera: Nymphalidae) based on five gene regions. Mol Phylogenet Evol 53, 463–478 doi:10.1016/j.ympev.2009.06.021 (2009).

84 Lanfear, R., Hua, X. & Warren, D. L. Estimating the Effective Sample Size of Tree Topologies from Bayesian Phylogenetic Analyses. Genome Biol Evol 8, 2319–2332 doi:10.1093/gbe/evw171 (2016).

85 Lanfear, R., Calcott, B., Ho, S. Y. W. & Guindon, S. PartitionFinder: Combined Selection of Partitioning Schemes and Substitution Models for Phylogenetic Analyses. Mol Biol Evol 29, 1695–1701 doi:10.1093/molbev/mss020 (2012).

86 Drummond, A. J., Suchard, M. A., Xie, D. & Rambaut, A. Bayesian Phylogenetics with BEAUti and the BEAST 1.7. Mol Biol Evol 29, 1969–1973 doi:10.1093/molbev/mss075 (2012).

87 Rambaut, A., Drummond, A. J., Xie, D., Baele, G. & Suchard, M. A. Posterior Summarization in Bayesian Phylogenetics Using Tracer 1.7. Syst Biol 67, 901–904 doi:10.1093/sysbio/syy032 (2018).

88 Päivinen, J. et al. Negative density-distribution relationship in butterflies. BMC Biol 3, 5, doi:10.1186/1741-7007-3-5 (2005).

89 Paradis, E., Claude, J. & Strimmer, K. APE: Analyses of Phylogenetics and Evolution in R language. Bioinformatics 20, 289–290 doi:10.1093/bioinformatics/btg412 (2004).

90 R: A language and environment for statistical computing. https://www.R-project.org/ (R Foundation for Statistical Computing, Vienna, Austria, 2018).

